# Summary Visualisations of Gene Ontology Terms with GO-Figure!

**DOI:** 10.1101/2020.12.02.408534

**Authors:** Maarten JMF Reijnders, Robert M Waterhouse

**Author notes:** **Correspondence:** (MJMFR), (RMW).

## Abstract

The Gene Ontology (GO) is a cornerstone of functional genomics research that drives discoveries through knowledge-informed computational analysis of biological data from large-scale assays. Key to this success is how the GO can be used to support hypotheses or conclusions about the biology or evolution of a study system by identifying annotated functions that are overrepresented in subsets of genes of interest. Graphical visualisations of such GO term enrichment results are critical to aid interpretation and avoid biases by presenting researchers with intuitive visual data summaries. Amongst current visualisation tools and resources there is a lack of standalone open-source software solutions that facilitate systematic comparisons of multiple lists of GO terms. To address this we developed GO-Figure!, an open-source Python software for producing user-customisable semantic similarity scatterplots of redundancy-reduced GO term lists. The lists are simplified by grouping together GO terms with similar functions using their quantified information contents and semantic similarities, with user-control over grouping thresholds. Representatives are then selected for plotting in two-dimensional semantic space where similar GO terms are placed closer to each other on the scatterplot, with an array of user-customisable graphical attributes. GO-Figure! offers a simple solution for command-line plotting of informative summary visualisations of lists of GO terms, designed to support exploratory data analyses and multiple dataset comparisons.

## Introduction

Since its foundation more than two decades ago, the Gene Ontology (GO) knowledgebase has developed into the world’s gold-standard for comprehensively describing gene functions (Ashburner *et al.*, 2000; The Gene Ontology Consortium, 2019). GO terms capture biological knowledge in three formalised ontologies (Biological Process, Molecular Function, and Cellular Component) that are hierarchical with ‘child’ terms being more specific than their ‘parent’ terms. Being both human-readable and machine-readable, and capturing these relationships in a directed acyclic graph (DAG), make the GO a powerful tool for computational analyses of results from large-scale molecular biology experiments and genome-wide assays (Dessimoz and Škunca, 2017). Typically, these results identify subsets of genes of interest, *e.g.* with increased or decreased expression levels under different conditions or treatments, or with high or low population-level genetic variation. GO enrichment analysis is subsequently performed to determine what is special or different about the subset of genes with respect to their associated processes, functions, or components. These results can then be used to support or refute hypotheses, inferences, or conclusions about the biology or evolution of the study system.

Enrichment analysis is generally performed by testing for statistical overrepresentation of GO terms annotated to genes from the subset of interest, using enrichment tools such as the TopGO (Alexa and Rahnenfuhrer, 2020) or GOStats (Falcon and Gentleman, 2007) R packages. Test results from various enrichment tools are usually presented as lists of GO terms with their associated probabilities (p-values), possibly also with the counts of annotated genes in the foreground (subset of interest), or ranks and p-values from different classes of statistical tests. While comprehensive, these lists can be difficult to interpret and may be subject to biases when attempting to summarise the key results, mainly because they do not present information on the relationships amongst the listed GO terms. Graphical visualisations of enrichment results have therefore been developed to aid interpretation and present visual summaries (Supek and Škunca, 2017). TopGO can provide views of how the most significant GO terms are distributed over the GO DAG, and results from TopGO and GOStats can be explored using R graphics functions (R Core Team, 2020). However, online web-server applications such as AmiGO (Carbon *et al.*, 2009), GOrilla (Eden *et al.*, 2009), and REVIGO (Supek *et al.*, 2011), are much more widely used for visualising results and producing summary figures for publications. Of these, REVIGO summaries are particularly popular as they simplify lists of GO terms by grouping together those with similar functions, using semantic similarities. Graphical summaries of the redundancy-reduced results can be produced as scatterplots that place more semantically similar GO terms closer to each other on the plot. Several options are provided that allow for interactive user- customisation of the graphics, and plotting R scripts can be downloaded for further customisation.

Despite the importance of using up-to-date versions of gene-term annotations and the core GO (Wadi *et al.*, 2016), web-server applications often do not keep pace with the evolution of the ontology and the annotations. This can lead to inconsistent or erroneous output summary visualisations generated from non-version-matched input lists of enriched GO terms. Additionally, web-server applications generally require input lists of GO terms to be individually uploaded by the user for each analysis they wish to conduct, making it difficult to perform systematic comparisons of multiple lists. Exploratory data analysis is greatly facilitated by graphical data summaries (Lee *et al.*, 2020), so being able to visually compare results from multiple lists is particularly important in this context. For example, using different software, parameters, normalisations, or filters to quantify differential gene expression will impact the results of gene set enrichment analyses. Alternatively, if gene lists are defined by clustering procedures, varying the input features, distance functions, or clustering algorithms will impact cluster membership and consequently the results of enrichment analyses. Beyond data exploration, visualising similarities or differences amongst enriched terms from multiple lists can support interpretations or conclusions drawn from comparing results from different tissues, life-stages, conditions, treatments, or even different species or populations.

To enable comparisons of graphical summaries of many lists of GO terms we developed GO-Figure!, an open-source Python software for producing user-customisable semantic similarity scatterplots. Using the latest core GO, or user-defined versions to match their enrichment analysis results, similarity scores are used to group redundant GO terms from which a representative is selected for plotting in two-dimensional semantic space, with user-control over grouping thresholds and scatterplot graphical attributes. GO-Figure! offers a simple solution for command-line plotting of informative data summary visualisations to support exploratory data analyses and multiple dataset comparisons.

## Materials and Methods

GO-Figure! is a Python software designed to facilitate the graphical visualisation of results from GO term enrichment analyses. Redundancies in the list of significantly enriched terms are minimised according to user-specified thresholds using pairwise semantic similarities to group similar terms together and subsequently select a representative for the group. These redundancy-reduced term lists are then visualised in two-dimensional semantic space where more functionally similar terms are placed closer to each other on the scatterplot.

### Redundancy-Reduced GO Term Lists

Redundancy reduction aims to identify similar GO terms and group them together in order to reduce the total number of terms and simplify the summary visualisation. For a user-provided list of GO terms derived from the results of a GO term enrichment analysis such as those produced by TopGO or GOStats, pairwise semantic similarities are calculated with the commonly-used formula proposed by Lin (Lin, 1998):

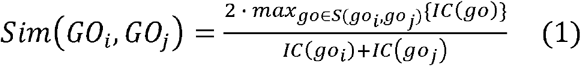

Here, *S* is the subset of GO terms shared between terms *i* and *j* after propagating up the GO DAG using the ‘is_a’ and ‘part_of’ relations. Information content (*IC*) is the relative frequency of a GO term *i* compared to the total number of GO terms in the UniProt (The UniProt Consortium, 2019) Gene Ontology Annotation (GOA) database:

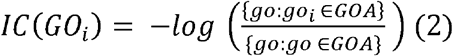

Each GO term is grouped with its most similar term if their pairwise semantic similarity score is higher than or equal to a user-adjustable threshold value. One GO term is then selected as the representative of the group following the logical reasoning outlined below. If the most similar term is already part of another group, then the GO term is added to this group and a representative term is selected between the newly added GO term or the current representative of the group. This process is repeated for all GO terms provided in the list from the enrichment analysis, resulting in groups of GO terms represented by a single term, based on their semantic similarities, with user-control over the resolution of redundancy reduction. GO terms with no other term above the semantic similarity score threshold remain as singletons on the scatterplot. Representative selection is based on the following stepwise logical reasoning:

1. If one GO term is annotated to 5% or more of the proteins in the GOA database (*i.e.* it is a term with a broad interpretation), select the other more specific GO term. If both terms are annotated to 5% or more, proceed to step 4.
2. If their p-values are substantially different, *i.e.* one GO term has a 50% higher p-value than the other, then select the term with the lower p-value.
3. If one GO term is a parent term of the other GO term, select the parent term.
4. If none of the above is true, select the first GO term as the representative (Python ordering of list objects results in a deterministic choice and therefore reproducible selections).

### Semantic Similarity Scatterplots

The redundancy-reduced lists of GO terms are summarised as circles plotted in two-dimensional semantic similarity space for each representative GO term, where more similar GO terms are plotted closer together on the x and y axes of the scatterplot (labelled semantic space X and Y). The pairwise semantic similarities for all representatives are used to generate the two-dimensional transformation using the multidimensional scaling algorithm from SciKit-Learn (Pedregosa *et al.*, 2011). By default, the colour of the point indicates the user-provided significance (log10 p-value) of that GO term (the representative term), and the size indicates the numbers of GO terms being represented by that GO term (the number of GO terms in the group). Several options are provided to allow users to easily configure how colours and sizes are used in plotting. The scatterplot itself is generated using the Seaborn (seaborn: statistical data visualization — seaborn 0.11.0 documentation) and Matplotlib (Hunter, 2007) Python packages.

### Gene Ontology Datasets

The calculations of semantic similarities and information contents detailed above rely on GO datasets including the core ontology from the GO Knowledgebase in Open Biomedical Ontologies (OBO) format that includes all parent-child relationships, and the GOA database of GO annotations from UniProt. Versions used to produce the figures in this manuscript: go.obo data version 18-11-2020, goa_uniprot_all.gaf version 10-7-2020. The importance of using up-to-date and version-matched gene-term annotations and the core GO used for enrichment analyses is often overlooked (Wadi *et al.*, 2016). GO-Figure! is provided with up-to-date go.obo ontologies and the precomputed GO term relations and information contents scores. User-provided input lists are checked for obsolete GO terms, which are updated if replacements are defined in the go.obo or discarded, with user notifications printed to a log file. As the core GO and the UniProt GOA database are regularly updated, support scripts and instructions are provided to compute the required GO term relations and information contents scores for user provided versions of the core GO and the UniProt GOA. To ensure that all GO terms present in the user-provided list can be assessed, it is recommended to use the same go.obo data version that was used for the GO term enrichment analysis, and the latest corresponding release of the GOA from UniProt.

## Results

### Summary Visualisations With GO-Figure!

GO-Figure! is designed to respond to two major gaps amongst current tools and resources for building summary visualisations of lists of GO terms: (1) a lack of GO-version-aware standalone software solutions; and (2) a lack of solutions that facilitate systematic comparisons of multiple lists. It is provided as an open-source Python software, which is simple to run even for novice Python users and requires no additional programming skills unlike other software that require running R packages or using application programming interfaces. It checks the validity of user-supplied lists with the latest ontologies and annotations, and provides user-control over version matching. As a standalone software it also allows for the rapid production of graphical summaries for multiple datasets without the need to repeatedly upload GO term lists to an online web-server. Summary visualisations are presented as scatterplots across two-dimensional semantic similarity space (see Materials and Methods), an intuitive data summary format popularised by REVIGO. Before plotting, semantic similarities and information contents are used to group similar GO terms and select a representative for each group, thereby achieving reduced redundancy of the displayed GO terms.

On the scatterplot, GO terms are arranged such that those which are most similar in semantic space X and Y are placed nearest to each other (Figure 1). By default, the p-value obtained from the enrichment analysis that generated the list is visualised using a gradient colour palette, and the sizes of the plotted points are scaled by the number of GO terms they represent. Figure 1A illustrates the basic functionality of GO-Figure! using an input list of 46 GO terms (biological processes) derived from data from (Van’t Veer *et al.*, 2002). This list is provided as an example dataset on the REVIGO web-server generated by analysing differentially expressed genes between breast cancer patients with fewer than five years to metastases and at least five years disease-free. Figure 1B shows the data summary produced selecting the Lin semantic similarity measure and otherwise using default settings on the REVIGO web-server for the same list of 46 GO terms.

**Figure 1.**
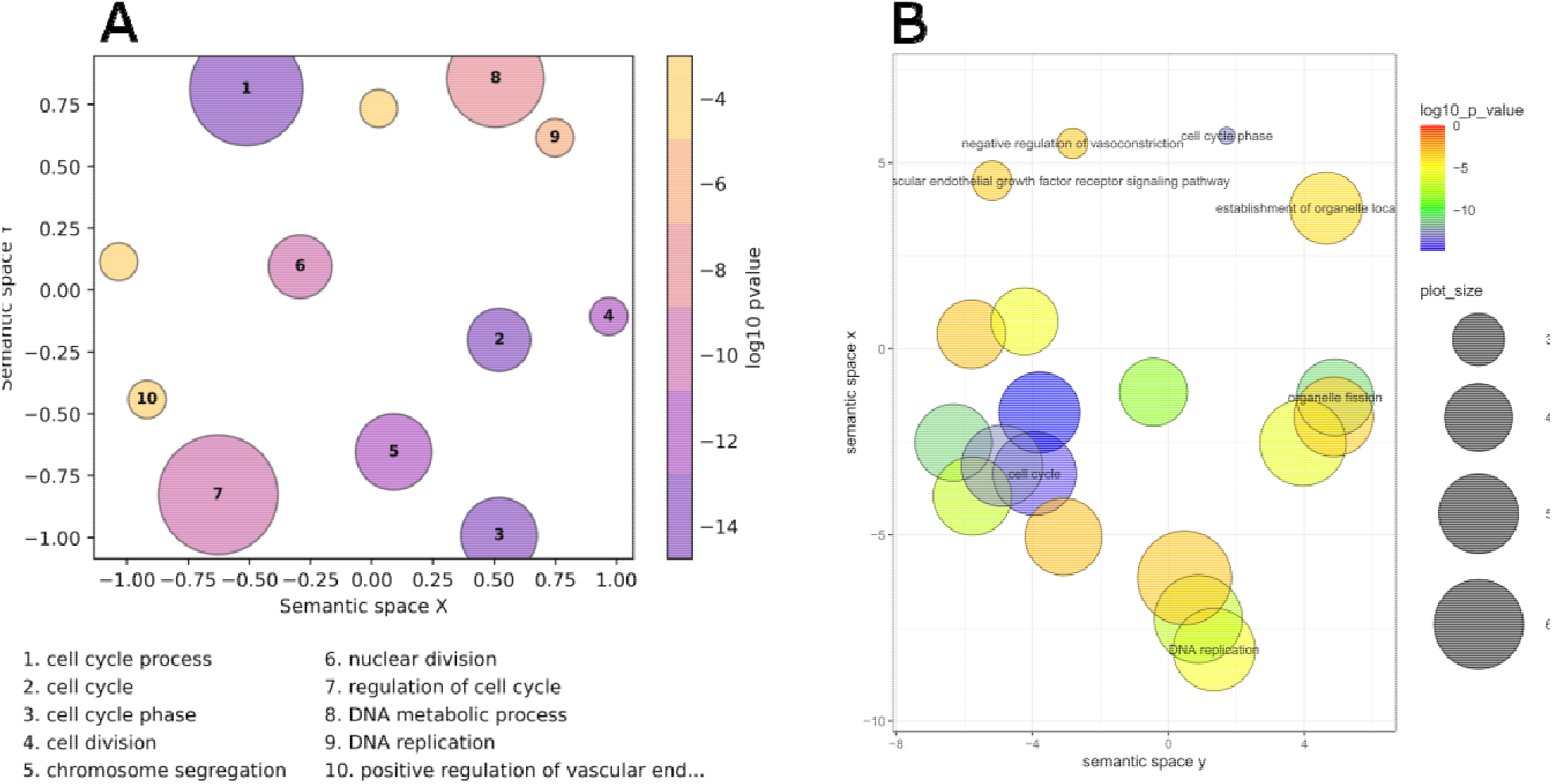
Semantic similarity scatterplots summarising a list of enriched Gene Ontology (GO) terms from a breast cancer differential gene expression study. The scatterplots show GO terms as circles arranged such that those which are most similar in semantic space X and Y are placed nearest to each other. They show results employing default plotting options with (A) GO-Figure! generated using the Seaborn and Matplotlib Python packages, and (B) REVIGO generated with the R script from the REVIGO web-server using the ggplot2 and scales R packages. Both use Lin (Lin, 1998) semantic similarities with a cutoff of 0.7 for redundancy reduction. The input GO term list from the REVIGO website consists of 46 Biological Process terms from an enrichment analysis of data from (Van’t Veer et al., 2002). Both scatterplots show the significance obtained from the enrichment analysis using a gradient colour palette (log10 p-value), and the sizes of the plotted circles are scaled by the number of GO terms they represent. In (A) the ten most significant terms are labelled with numbers and their descriptions are provided below the plot, in (B) the seven most significant terms are labelled with their descriptions.

GO-Figure! reduces redundancy to 12 representative GO terms, the ten most significant are labelled on the scatterplot and their descriptions are shown below (Figure 1A). Six are singletons and the other six represent the remaining 40 GO terms, the smallest group two (*organelle fission* and *nuclear division*) or three (*chromosome segregation*, *sister chromatid segregation*, and *mitotic sister chromatid segregation*) similar terms, and the largest groups 14 terms represented by *regulation of cell cycle* (Table S1). GO-Figure! recognises one term identifier (GO:0007067) as obsolete and instead uses the replacement term identifier (GO:0000278) as defined by the latest core GO. REVIGO only reduces redundancy to 19 representative GO terms, seven of which are labelled with their descriptions on the scatterplot (Figure 1B), employing the January 2017 version of the GO and the March 2017 version of the UniProt GOA. Singletons make up 12 of the representatives and the remaining 34 GO terms are represented by seven groups, the smallest groups two (*cytoskeleton organization* and *DNA conformation change*) similar terms, and the largest groups 14 terms represented by *cell cycle process* (Table S2).

### Customising Summary Visualisations

GO-Figure! allows for user-customisation to tailor three main features of the resulting scatterplots: (1) colour palettes and scaling; (2) management of GO term labels; and (3) incorporating user-provided additional data. This is achieved through a wide selection of command line options detailed in the software help and the user guide. A large choice of colour schemes is made available through the full library of colour brewer palettes provided by the MatPlotLib Python package, *e.g.* the popular plasma or viridis palettes (Figure 2). By default, as shown in Figure 2A, circles are coloured based on the user-provided p-values (log10-scaled) for each GO term, and size-scaled by the number of GO terms they represent (members, resulting from the redundancy reduction steps). Colours can instead be used to indicate the number of members, the frequency of the GO term in the GOA database (information content score), or simply a unique colour per GO term. Similarly, p-values, members, or GOA frequency, can be used to size-scale the points, or a fixed size can be applied.

**Figure 2.**
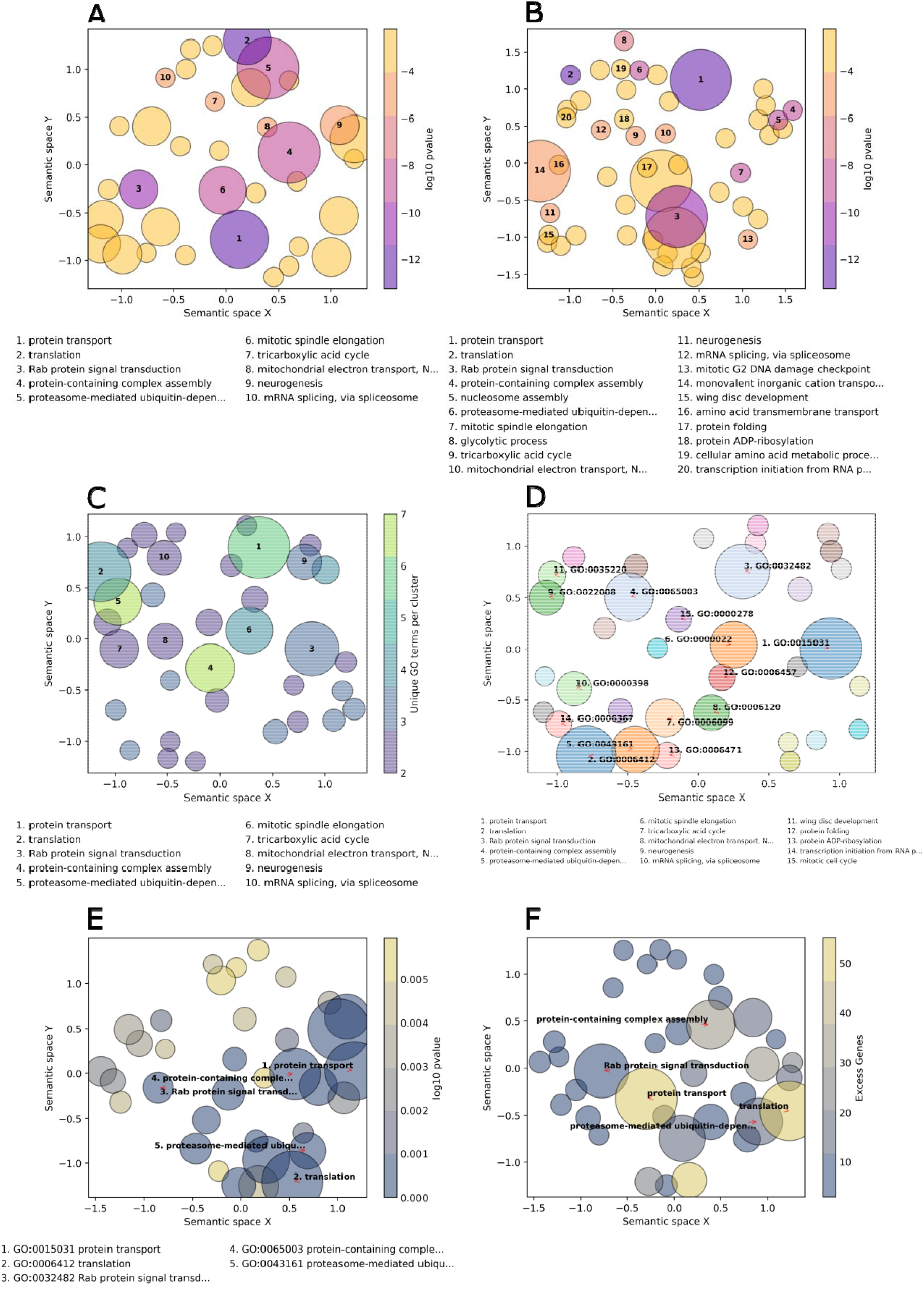
Example customisations of GO-Figure! summary visualisations. The input GO term list consists of 60 Biological Process terms from an enrichment analysis of genes with low protein sequence divergence rates from (Neafsey et al., 2015). (A) Default plotting options, i.e. semantic similarity threshold of 0.5 and ten numbered labels. (B) Increasing the semantic similarity threshold to 0.8 and plotting labels for the 20 most significant GO terms. (C) Using the colour palette ‘viridis’, with semantic similarity threshold of 0.5, colours based on the number of GO terms being represented (members), and sizes based on p-values. (D) Colour palette ‘tab20’ with a unique colour per circle, semantic similarity threshold of 0.5, three- column reduced font size legend for 15 GO term identifier labels with arrows. (E) Colour palette ‘cividis’ with colours based on p-values and sizes based on user-provided values, five character-limited descriptor labels, with GO term identifiers and descriptors in the legend. (F) Palette ‘cividis’ with colours based on user-provided values (excess genes) and sizes based on p-values, five descriptor labels with no legend.

The number of GO terms to be plotted depends on how many were supplied in the input list and on the stringency applied during redundancy reduction. To avoid cluttering the scatterplot with GO term descriptions, the most significant terms are labelled with numbers and their descriptions are printed below the scatterplot (Figure 2A-C). Command line options allow users to determine the numbers of GO terms to be labelled and manage the layouts of the descriptions (font sizes, number of columns, maximum character lengths of descriptors, etc.). The GO terms on the scatterplot can instead be labelled with their identifiers, or with their descriptors, using automatic label placement (Flyamer *et al.*, 2020) to optimise readability (Figure 2D-F). This provides the user with the flexibility to easily increase or decrease the amount of in-plot labelling text in a standardised manner.

The minimal user-provided input information required for building the summary visualisations is the input list of GO term identifiers with their corresponding probabilities resulting from enrichment testing (standard input). GO-Figure! is also able to process default files produced from enrichment testing with TopGO or GOStats, consisting of seven columns as detailed in the user guide. The values from the ‘Significant’ (TopGO) and ‘Count’ (GOStats) columns, which provide counts of the numbers of genes in the foreground set annotated with the GO term, can be used to determine plotting colours and sizes. Alternatively, users can prepare a standard-plus input consisting of the GO term identifiers, their corresponding probabilities, and a third column with a user-defined metric that can also be used to determine plotting colours and sizes (Figure 2E-F). For example, the total numbers of genes annotated with the GO term in the specific annotation dataset used during enrichment testing, rather than frequencies calculated from the entire UniProt GOA database.

The scatterplots aim to summarise information through redundancy reduction, hence only representative GO terms are shown for groups of similar GO terms. For reference, group membership information is provided in tab-delimited text files as part of the default output. In addition, for tracking provenance and facilitating reproducibility a log file is created to record the versions of GO-Figure!, the core GO, and UniProt GOA database used, as well as the options and parameters used to produce the scatterplot, and to track any GO terms that were either updated to correspond with the current version of the GO or that were discarded because they were found to be obsolete with no replacements. Finally, the Python data frames used to create the scatterplots are also saved as tab-delimited text files. This allows for users with more advanced experience of MatPlotLib functions to further customise the summary visualisations, to integrate them into multi-panel figures, or to create interactive versions of the scatterplots.

## Conclusion

Recognising the usefulness and popularity of semantic-similarity-based redundancy reduction for producing visualisations of GO term lists, GO-Figure! offers a simple solution for command-line plotting of informative graphical summaries. It addresses a lack amongst current visualisation tools and resources of standalone open-source software solutions that are GO-version-aware and facilitate systematic comparisons of multiple lists of GO terms. Summary visualisations of gene set functional annotations with GO-Figure! enable researchers to perform exploratory data analyses and multiple dataset comparisons to support hypotheses or conclusions about the biology or evolution of their study systems.

## Availability

GO-Figure! is available as a set of python scripts with associated datasets. Instructions on the installation, updatability, and use of GO-Figure! are available from https://gitlab.com/evogenlab/GO-Figure

## Author Contributions

Conception (MJMFR and RMW). Software development (MJMFR). MJMFR and RMW wrote the manuscript. All authors read and approved the manuscript.

## Acknowledgements

The authors thank Romain Feron, Livio Ruzzante, and Antonin Thiébaut for beta-testing the software and useful suggestions for improvement, as well as for feedback on the manuscript.

## Supplementary Materials

Supplementary Tables S1 and S2 are provides as an Excel file: Supplementary_Tables_S1_S2.xlsx

**Table S1**

GO-Figure! summary data of input GO terms and their representatives after redundancy reduction as shown in Figure 1A. For each GO term columns present: representative GO term, member GO term, GO term description, p-value from enrichment testing, computed information content (IC), and computed frequency in the UniProt GOA.

**Table S2**

REVIGO summary data of input GO terms after redundancy reduction as obtained from the REVIGO web-server to produce Figure 1B, with the addition of a column to indicate the representative GO term selected during redundancy reduction.

